# Molecular genetic diversity of donkey (*Equus asinus*) in South Korea

**DOI:** 10.1101/213520

**Authors:** Sungwook Yun, Giljae Cho

## Abstract

The research on domestic donkey has little information and criteria comparing to other livestock, companion animal in South Korea. We analyzed genetic database of domestic donkey using microsatellite marker to clarify domestic donkey identification and paternity test. It is the first microsatellite marker analysis on domestic donkey in South Korea. We studied 179 horse samples from 50 Thoroughbred, 50 Jeju Halla horse, including 79 donkeys then analyzed 15 microsatellite marker (AHT4, AHT5, ASB2, ASB17, ASB23, CA425, HMS1, HMS2, HMS3, HMS6, HMS7, HTG4, HTG10, LEX3 and VHL20). We observed genetic diversity from biostatic analysis of them.

The number of alleles on total average is 6.08 observed from 1 (ASB17), 2 (HMS1) to 14 (AHT5). The observed heterozygosity (OHet) is from 0.0000 (ASB17, HMS1) to 0.8608 (ASB23) which is mean value of 0.4861, the expected heterozygosity (EHet) is from 0.0000 (CA425) to 0.9104 (AHT5) with mean value of 0.5915, and the Polymorphism Information Content (PIC) on each group of microsatellite marker is from 0.0000 (ASB17) to 0.8968 (AHT5) observed as a mean value of 0.5374. Among 15 markers AHT4, AHT5, ASB23, CA425, HMS2, HMS3, HTG4, HTG10, LEX3 is observed above 5.000.

The results on 15 microsatellite marker analysis of 3 horse groups is that the donkey had 0.5915 EHet, 0.4861 OHet on average, Thoroughbred had 0.6721 EHet, 0.6587 OHet on average, and the Jeju halla horse had 0.7898 EHet, 0.7093 OHet on average observed. Furthermore the mean alleles value is observed as 6.08, 4.83, 8.00 in donkey, Thoroughbred, Jeju halla horse breed in each.

## 1. Introduction

Donkey is a domesticated animal belonging to Equidae along with horse and zebra. It is assumed that the domestication of *Equus asinus* took place in 6000 BC in North Africa (Egypt area) from Nubian wild ass and Somalian wild ass [2, 21]. Donkey has worked with mankind for centuries, and the most important role was transportation. Donkey still remains an important work stock animal in poor territories.

The height of donkey’s withers is very diverse depending on the species, known to be average 102 cm. The coat color, too, varies, such as white, gray and black and it has a black stripe from the mane to the tail and another stripe sideways on the shoulders. The mane is short and straight up, and the shape of the tail with long hairs only at the tip is closer to a cow than a horse. The ears are very long and the base and tip are black. Unlike a horse, donkey’s chestnuts are located only on the front legs.

It is assumed that domestically bred donkey has been introduced from northern regions such as China and Mongolia, and it is assessed that currently, in South Korea, about 800 horses of donkey are bred, but the exact breed, introduction route and breeding scale are not well known. According to the study of Yun [27], it was observed that in the domestically bred donkey, the ear length was 17~28 cm (average 23 cm); withers height, 90~135 cm (average 118.3 cm); and body length, 109~150 cm (average 131.2 cm). It was reported that the distribution of coat colors was white, chestnut, gray, black and brown. The domestic feeding status is that they are bred mostly in Gyeonggi-do, Jella-do, Gyeongsang-do and Gangwon-do regions.

The purposes of breeding are mostly for experience, tourist carriage, or eating, but there are no basic data for the utilization of domestically bred donkey and the preservation of the blood. A DNA marker refers to short DNA fragment with polymorphism on DNA sequence that can indicate the location of a specific gene on the chromosome. For various genes and DNA markers involved in an economic trait, when DNA markers and these character genes are located close to each other, according to the degree of genetic recombination, the basis of Mendelian genetics, there are regions of the genes in which almost no gene cross occurs among them. Thus, the gene map that indicates the locations of specific DNA markers and these markers is the starting point and the key tool [3].

Recently, in most animals like a man, the DNA analysis technique using microsatellite markers has been applied to the differentiation of individual, paternity test, the preservation of endangered animals, phylogenesis related to the origin and production traceability.

It is known that generally, DNA constitutes chromosome with functional DNA, which includes genetic information and nonfunctional DNA, which does not include genetic information. In the DNA, which does not include genetic information, there is a tandem repeat sequence region, and the repeating unit of this region formed by the repetition of sequences of 10-50 bases is called Variable Number of Tandem Repeat (VNTR) and the repeating unit formed by the repetition of sequences of 2~7 bases is called Short Tandem Repeat (STR) or Microsatellite [4]. Microsatellite is sized about 100~400 bp. Even in severely damaged DNA specimens, it can be amplified, and gene inspection can be conducted with a small amount of DNA while VNTR has a lot of allelic genes, so it is difficult to measure its exact size, but microsatellite has fewer number of allelic genes than VNTR, so it is possible to identify them accurately. Thus, microsatellite markers are widely used for the differentiation of individuals of animals, including human or paternity test [11, 26].

Recently, in most animals like a man, the DNA analysis technique using microsatellite markers has been applied to the differentiation of individual, paternity test, the preservation of endangered animals, phylogenesis related to the origin and production traceability. In addition, countries throughout the world have widely used microsatellite markers since the mid-1990s for the purposes of the hereditary diversity of the country’s traditional domestic animals, origination and system, hereditary characteristics and preservation [1, 5].

Microsatellite refers to numerous repeats of simple base sequences existing in genomes of an organism, which is abundantly distributed throughout the entire spinal animal genomes in short, simple and repetitive base forms [12].

Microsatellite has a high mutation rate over 1/10⁴~ 1/10⁶ per generation and has high viscosity according to the group. These characteristics show specific polymorphism at the object level as well as in the group, so it is used as a useful tool for genetic mapping as well as information about the hereditary diversity of animals, including human and plants [8, 10, 13, 14, 24].

Polymorphism of microsatellite is formed by the number of repeats of the basic repeating unit, which is inherited according to Mendel’s law of inheritance of a half from parents. Microsatellite locus evenly distributed in chromosome DNA shows various variations between individuals, so it is used for an analysis of the relationship between individual breeds and as an important marker for chromosome mapping. Till now, nothing has been known about the function of microsatellite, but it was reported that it is the region appearing common in the genes of all vertebrates, the length of base sequence varies depending on the number of repetitions, and each marker has many allelic genes. This length polymorphism is inherited to offspring from the mother and father according to Mendel’s law, existing in the form of allelic genes on the chromosome and showing hereditary polymorphism by each individual [14]. In parentage diagnosis, microsatellite and minisatellite can be used simultaneously, but using microsatellite used as a standard for international genetic mapping can easily secure amplified primer, observe various genotypes and check the unique hereditary characteristics of the individual, so it is usefully utilized as a marker for the blood registration through the differentiation of individuals and parentage diagnosis [13, 24]. Microsatellite in a horse was reported for the first time by Ellegren et al. [8] and Marklund et al. [14].

It is judged that in South Korea, donkey will be in the limelight as a high-value genetic resource in the future. In addition, it may be disguised as horse meat. To monitor this, it is necessary to protect breeding farms through a donkey meat traceability system. This study aims to secure basic data to increase the protection and utilization of genetic resources of domestic donkey, getting ready for the traceability system of domestic donkey using microsatellite markers and investigating the hereditary diversity of donkey.

## 2. Materials and Methods

### 2.1. Sample collection and DNA extraction

Genomic DNAs from whole blood samples of 179 horse breeds [79 donkey, 50 Thoroughbred and Jeju Halla horse (Thoroughbred and Jeju horse crossbred)] were extracted using a MagExtractor System MFX-2000 (Toyobo, Osaka, Japan) according to the manufacturer's protocols [24].

For the quantification of the separated DNA, the absorbance was measured at the 260 nm and 280 nm wavelengths using Nanodrop^TM^ 8000 Spectrophotometer (Thermo, USA), and DNA extracted based on absorbance at 260 nm at the value of 1.0 (Path length = 10.0 mm) was diluted, and the concentration was adjusted to 50 ng/ul. In addition, samples with too high or too low A260/A280 ratio based on 1.8 purity was judged to be low, and DNA was re-extracted from the blood to use in the experiment.

To separate and check the quantified DNA by the naked eye, the finally extracted DNA was checked by electrophoresis on 2.5% Agarose at 100V for 30 min. using Mupid-2 Plus Electrophoresis Cell (TaKara, Japan).

### 2.2. Microsatellite markers and analysis

Fifteen microsatellite markers (AHT4, AHT5, ASB2, ASB17, ASB23, CA425, HMS1, HMS2, HMS3, HMS6, HMS7, HTG4, HTG10, LEX3 and VHL20) were used for analysis of the horse breeds. PCR was performed according to the manufacturer’s protocols. Of the 15 markers, with markers ASB17, ASB23, CA425, HMS1 and LEX3, a single PCR was conducted. As for composition for PCR, template DNA 2 μl, 10 Pmol forward and reverse primer 2 ul, respectively and sterile distilled water 2 ul were mixed on PCR Premix buffer (Qiagen, Germany), adjusted to 15 μl in total, and then, amplified by GeneAmp PCR system 9700 (Applied Biosystems, USA).

In the PCR process, heating at 95°C for 10 min. to induce degeneration and three steps of denaturation at 95°C for 30 sec., annealing at 60°C for 30 sec., and extension at 72°C for 60 sec. were repeated 30 times in total, and lastly, the final extension process was made at 72°C for 60 min.

For the single PCR, template DNA 2 μl, 10 Pmol forward and reverse primer 2 ul, respectively and sterile distilled water 6.5 ul were mixed on PCR Premix buffer (Qiagen, Germany), adjusted to 25 μl in total, and then, amplified by GeneAmp PCR system 9700 (Applied Biosystems, USA).

In the PCR process, heating at 95°C for 5 min. to induce degeneration and three steps of denaturation at 95°C for 30 sec., annealing at 60°C for 30 sec., and extension at 72°C for 60 sec. were repeated 35 times in total, and lastly, the final extension process was made at 72°C for 60 min.

With the amplified DNA, dielectrolysis was made with 2.5% agarose gel, and by comparing the amplification and concentration indirectly, it was tested for genotype determination.

The genotype analysis after the PCR was as follows: Mixing the amplified fragment 0.5 ul, Gene Scan 500 RIZ size standard (Applied Biosystems, USA) 0.25 ul and deionized Hi-Di formamide (Applied Biosystems, USA) 12.25 ul well to make the final volume 13 ul; denaturation at 95°C for three min. and dipping on ice for three min.; loading the denatured PCR product on automatic gene analyzer (ABI 3130 xl Genetic Analyzer, USA); and electrophoresis on POP 7 polymer (Applied Biosystems, USA) at 15 kV. Then, with the peak row data, the size of allelic genes (base pair) for each marker was determined based on the result of 2015/2016 Horse Comparison Test No. 1 of the International Society for Animal Genetics (ISAG) using GeneMap Software ver. 4.0 (Applied Biosystems, USA).

### 2.3. Statistical analysis

Analysis of the hereditary diversity of domestically bred *Equus asinus*, that is, the observed heterozygosity (OHet), expected heterozygosity (EHet), number of allelic genes and frequency was conducted, using Microsatellite Marker Tool Kit Ver. 3.1.1 (Microsoft®, USA) [19], and the Polymorphism Information Content (PIC) of the groups for each microsatellite marker analyzed [6] was calculated through the methods of Marshall et al. [15] and Nei et al. [16].

To examine the cousin relation of domestically bred *Equus asinus*, for the estimation of DA genetic distances for an analysis of the relationships among the groups, the distances were calculated, using DISPAN Package [18], a population genetics analysis program that uses the method of Nei et al. [16], and using the DISPAN, a phylogenetic tree was drawn up based on the hereditary distances between the groups, through Unweighted Pair-group Method With Arithmetic Average (UPGMA) [22] method. Based on the estimated value of the individual hereditary distances of all groups, for drawing up DA genetic distances among all individuals, Phylip Ver.3.69 statistics program, a population genetics analysis program was used based on the frequency of individual allelic genes through the level of simple allele-sharing measurement [9].

## 3. Results

### 3.1. Analysis of the genetic diversity of donkey

As shown in Table 1, it was observed that the number of allelic genes was 1 (ASB17) and 2 (HMS1) to 14 (AHT5), 6.00 on average. The OHet was 0.0000 (ASB17, HMS1) to 0.8608 (ASB23), 0.4861 on average. The EHet was 0.0000 (CA425) to 0.9104 (AHT5), 0.5915 on average; and the Polymorphism Information Content (PIC) of groups by each microsatellite marker was 0.0000 (ASB17) to 0.8968 (AHT5), 0.5374 on average. Of the 15 markers, in AHT4, AHT5, ASB23, CA425, HMS2, HMS3, HTG4, HTG10, LEX3, it was higher than 5.000.

**Table 1.**
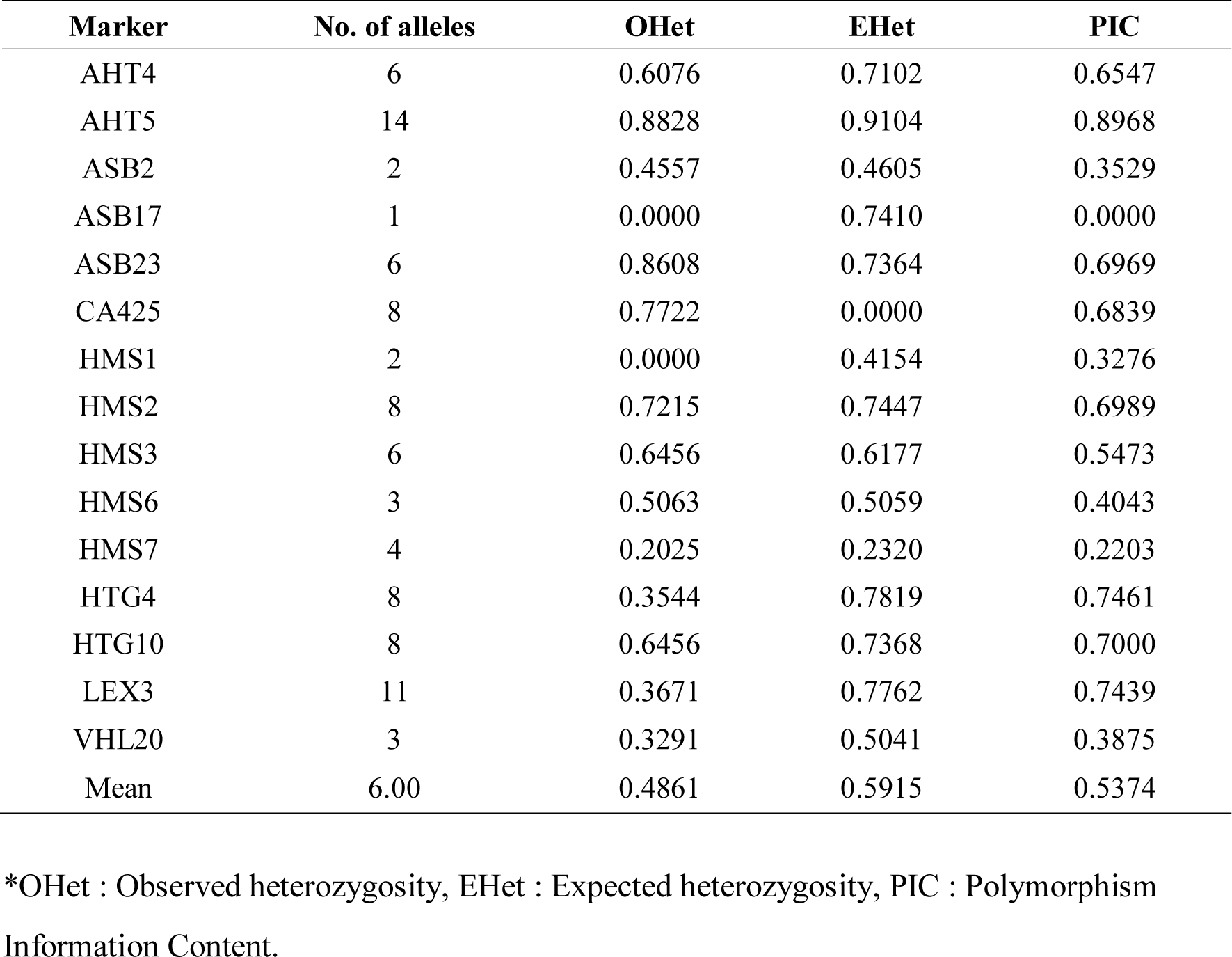
Number of allele, heterozygosity, and PIC of the 15 microsatellite markers in 79 donkeys

As a results of an analysis of microsatellite DNA types of donkey, Thoroughbred and Jeju Halla horse, allelic genes and frequency are as shown in Table 2 and 3. As a result of an analysis of 15 microsatellite markers with the three horse groups, the average Ehet and Ohet were 0.5915 and 0.4861, respectively in donkey; 0.6721 and 0.6587 in Thoroughbred; and 0.7898 and 0.7093 in Jeju Halla horse. In addition, it was observed that the average number of allelic genes was 6.00, 4.83 and 8.00, respectively in donkey, Thoroughbred and Jeju Halla horse.

**Table 2.**
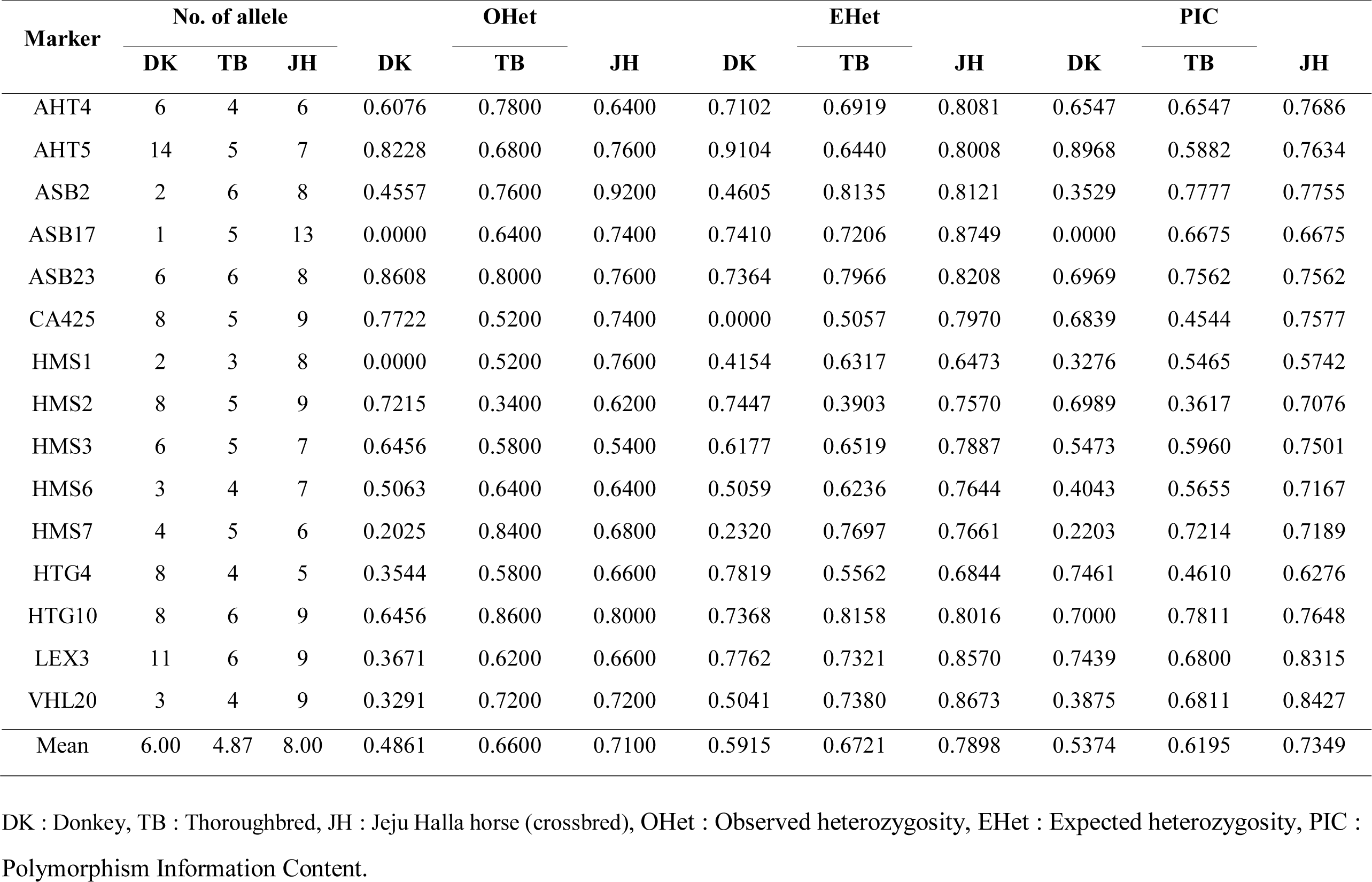
Number of allele, heterozygosity, and PIC of the 15 microsatellite markers in 179 horse breeds

**Table 3.** Expected, Observed heterozygosity and mean number of alleles(MNA) observed across 15 microsatellite loci for each population

| Population | Sample size | ±Unbiased HzSD | ±Obs HzSD | MNA | MNA SD |
| --- | --- | --- | --- | --- | --- |
| DK | 79 | $\pm 0.59150.0624$ | $\pm 0.48610.0145$ | 6.00 | 3.63 |
| TB | 50 | $\pm 0.67210.0310$ | $\pm 0.65870.0173$ | 4.83 | 1.03 |
| JH | 50 | $\pm 0.78980.0160$ | $\pm 0.70930.0166$ | 8.00 | 1.89 |

DK : Donkey, TB : Thoroughbred, JH : Jeju Halla horse (crossbred).

Based on the result of an analysis of microsatellite markers, dendrograms of groups for the standard genetic distance and the minimum genetic distance were drawn, using Unweighted

Pair Group Method with Arithmetic mean (UPGMA) and Neighbor Joining (NJ) clustering method, based on the genetic matrix and presented in Figure 1. To compare the dendrograms, in three breeds 179 horses, donkey and Thoroughbred breed formed a clearly different group, but it was observed that Jeju Halla horse formed a group, mixed with a Thoroughbred horse.

**Fig. 1.**
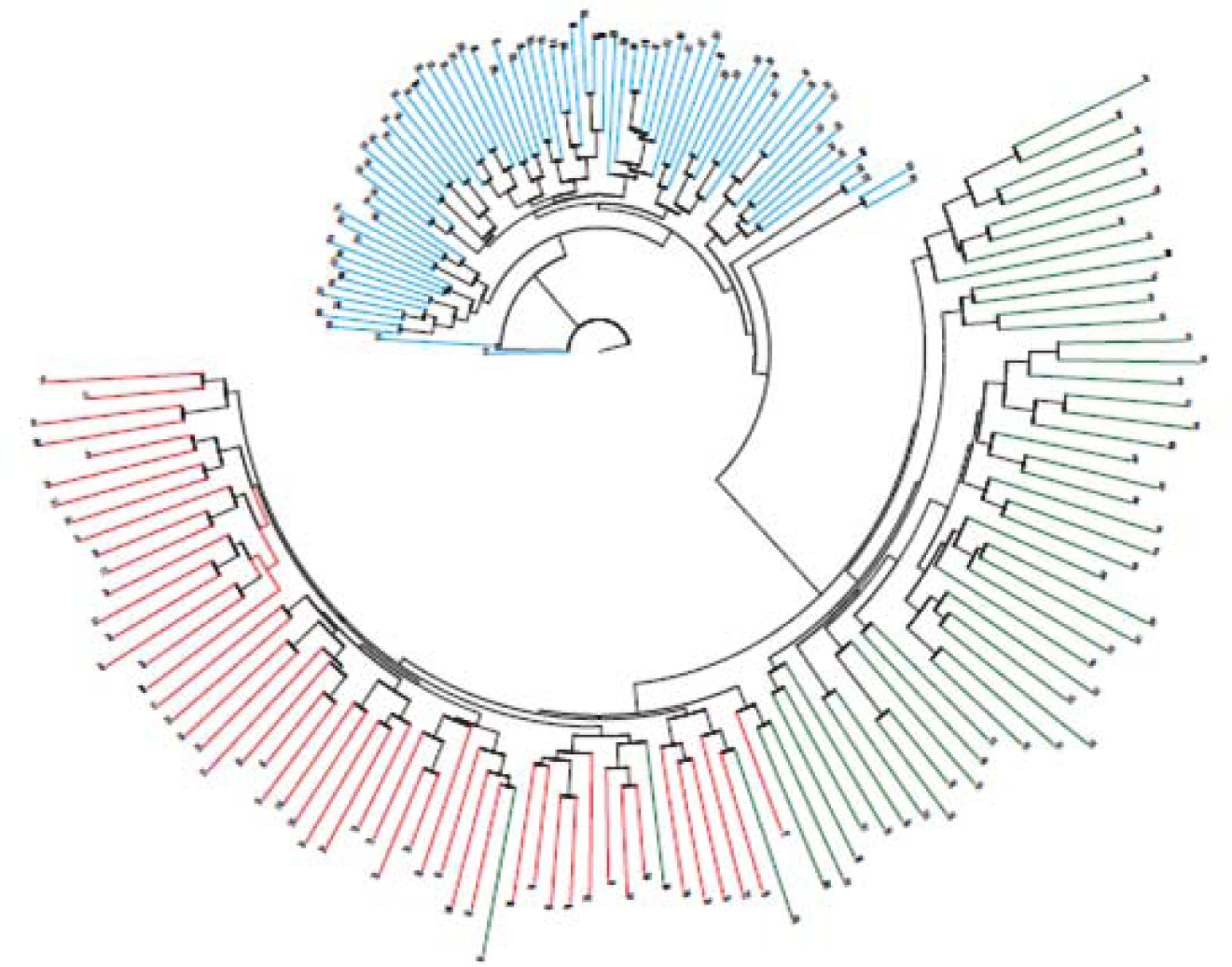
Phylogenetic tree showing allele-sharing distances among 179 individuals in 1 donkey and 2 horse breeds. Blue color: donkey, Green color: Thoroughbred, Red color: Jeju Halla horse.

## 4. Discussion

To meet the demands of the domestic donkey market and secure better quality donkey and donkey meats, most of all, it is necessary to select donkey with excellent blood and formulate and enhance donkey breeding technology. However, there are considerably very insufficient investigations of the breeding and genetic of donkey as compared to those of Thoroughbred, Jeju horse or Jeju Halla horse in South Korea.

Genotype is a stable unit at the object level but becomes an unstable unit through generations. In Mendel’s the law of segregation, “Genotype is the combination of two allelic genes, and one side of the allelic genes, only, is delivered to the next generation in the same probability,” the unit stably delivered, transcending generations of the components of genotype is not genotype but allelic genes [3].

In a shift in generations, that only one of the allelic genes is delivered at the same probability means that, since individual’s genotype is the combination of two allelic genes, the probability at which one allelic gene is delivered is 1/2. At the chromosome level, allelic genes are part of chromosome phase on one side, and in Molecular Biology, it is polymorphism such as SNP and VNTR on one chromosome. For example, in SNP, allelic gene becomes one side of the base of T or C [3, 23].

As compared to genotype frequency, gene frequency is very stable beyond the generation, so usually, many studies of molecular evolution deal with changes in gene frequency. However, what is directly related to the phenotype of individual is genotype [7, 20].

As a results of an analysis of genotype distribution using microsatellite DNA markers with domestically bred donkey to understand hereditary characteristics and a study to secure baseline data for the protection of donkey’s unique genetic resources and the promotion of high value added donkey, it was observed that the average EHet and OHet were 0.5915, 0.4861, respectively in domestically bred donkey; 0.6721, 0.6587 in Thoroughbred; 0.7898, 0.7093 in Jeju Halla horse. Also, it was observed that the average number of allelic genes was 6.00, 4.83 and 8.00 in donkey, Thoroughbred and Jeju Halla horse, respectively. It was noted that *Equus asinus* or Thoroughbred was fixed into a single breed. However, it is assumed that Jeju Halla horse had high OHet, EHet and number of allelic genes because they are cross-bred horses, in which the genes of various breeds were mixed. In addition, it was observed that microsatellite markers AHT4, AHT5, ASB23, CA425, HMS2, HMS3, HTG4, HTG10 and LEX3 were Polymorphism Information Content (PIC) over 5.000, so it is expected that they can be utilized in the differentiation of individuals of *Equus asinus* or paternity test. Based on PIC value of each marker, the validity and reliability of the marker can be estimated, and if PIC value is higher than 0.5000, it is judged that the reliability of the marker is valid for blood analysis. If it is higher than 0.7000, it is known that it has universal validity for analysis and can get a result of high reliability.

In a single gene locus, the indicator that expresses the diversity of a group is heterozygosity. Heterozygosity is defined as “the probability that, when randomly two allelic genes are extracted from a group, the two may differ.” In an association analysis or linkage disequilibrium analysis, to be used as a marker, the higher the heterozygosity, the more desirable it becomes [10, 21, 25]. Even when the plural groups are mixed, of course, heterozygosity increases, but if there is no mix of groups, generally, heterozygosity is related to mutation. The higher the mutation rate and the higher the effective size of the group, the greater the heterozygosity (the diversity of the group) becomes [17].

In general, in an analysis of genetic characteristics using a microsatellite marker, heterozygosity can be judged from the basic figure by which the degree of mixing of the target varieties and other varieties are predicted. In general, if pure blood is preserved through powerful selection without a mix of species from the outside, the value of heterozygosity appears low, and if there is a mix of different breeds, it is observed that the value of heterozygosity is high. However, since the value of heterozygosity becomes higher as there are more individuals used in the study, it is difficult to judge the mix of species only based on the value of heterozygosity. As a result of an analysis of three horse groups with 15 microsatellite markers, it was observed that the average EHet and OHet were 0.5915 and 0.4861 in donkey; 0.6721 and 0.6587 in Thoroughbred; and 0.7898 and 0.7093 in Jeju Halla horse. In addition, as a result of drawing up and analyzing a dendrogram of groups about the standard genetic distance and minimum genetic distance, in three breeds, 179 horses, donkey and Thoroughbred breed formed a clearly different group, but it was observed that Jeju Halla horse formed a group, mixed with Thoroughbred horse. Heterozygosity of domestically bred donkey was lower than two species and formed a group clearly differentiated like Thoroughbred horse. It is assumed that heterozygosity is low because of few breeding heads of domestically bred donkey and inbreeding by limited male horses.

### 4.1. Conclusion

We analyzed the first genetic database of domestic donkey using microsatellite marker to clarify domestic donkey identification and paternity test in South Korea. The donkey was observed that the number of allelic genes was 1 (ASB17) and 2 (HMS1) to 14 (AHT5), 6.00 on average. The OHet was 0.0000 (ASB17, HMS1) to 0.8608 (ASB23), 0.4861 on average. The EHet was 0.0000 (CA425) to 0.9104 (AHT5), 0.5915 on average. The PIC of groups by each microsatellite marker was 0.0000 (ASB17) to 0.8968 (AHT5), 0.5374 on average, and 9 microsatellite markers (AHT4, AHT5, ASB23, CA425, HMS2, HMS3, HTG4, HTG10 and LEX3) were higher than 5.000. These markers are useful

## Acknowledgement

This study was supported by Korea Institute of Planning and Evaluation for Technology in Food, Agriculture, Forestry and Fisheries in the field of business of technique in Agriculture and Bio industry (Project number: 316026).

We would like to thank Cho Cangyeon, Animal Genetic Resources Research Center, National Institute of Animal Science Rural Development Administration, Republic of Korea, for his comments on the Statistical analysis.

